# A computational approach to estimating nondisjunction frequency in *Saccharomyces cerevisiae*

**DOI:** 10.1101/024570

**Authors:** Daniel B. Chu, Sean M. Burgess

## Abstract

Errors segregating homologous chromosomes during meiosis result in the formation of aneuploid gametes and are the largest contributing factor to birth defects and spontaneous abortions in humans. *Saccharomyces cerevisiae* has long served as a model organism for studying the gene network supporting normal chromosome segregation. Current methods of measuring homolog nondisjunction frequencies are laborious and involve dissecting thousands of tetrads to detect missegregation of individually marked chromosomes. Here we describe a holistic computational approach to determine the relative contributions of meiosis I nondisjunction and random spore death in mutants with reduced spore viability. These values are based on best-fit distributions of 4, 3, 2, 1, and 0 viable-spore tetrads to observed distributions in mutant and wild-type strains. We show proof-of-principle using published data sets that the calculated average meiosis I nondisjunction frequency closely matches empirically determined values. This analysis also points to meiosis I nondisjunction as an intrinsic component of spore inviability in wild-type strains. We uncover two classes of mutants that show distinct relationships between nondisjunction death and random spore death. Class I mutants, including those with known defects in establishing and maintaining the physical engagement of homologous chromosomes display a 4-fold greater ratio of nondisjunction death to random spore death compared to Class II mutants, which include those with defects in sister chromatid cohesion. Low numbers of required tetrads facilitates epistasis analysis to probe genetic interactions. Finally the application of the R-Scripts does not require any special strain construction and can be applied to previously observed tetrad distributions.

## Introduction

Meiosis is an integral developmental program required for sexual reproduction in eukaryotes (PETRONCZKI *et al.* 2003). Through two rounds of chromosome segregation, the DNA content of parent diploid cells (2n) is reduced to form haploid gametes (1n). Homologous chromosomes separate from one another in the first meiotic division that follows meiosis I prophase. Proper separation requires a series of dynamic chromosome events that physically tether homologous chromosomes together to form a bivalent. In female mammals, meiosis I prophase occurs in ovaries of the fetus, yet cells are blocked from completing anaphase until sexual maturity (NAGAOKA *et al.* 2012). Thus, these contacts must be robust enough to last through decades spanning a reproductive lifespan.

Errors in chromosome segregation are the leading cause of birth defects in humans (HASSOLD AND HUNT 2001). Many instances can be traced to defects occurring during meiosis I prophase. Failure to properly separate homologs during meiosis I anaphase can result in chromosome aneuploidy in the fertilized zygote giving rise to live births, including trisomy 21, which is the cause of Down syndrome. A maternal age effect increases the incidence of meiosis I (MI) chromosome missegregation in these gametes with compromised homolog attachments. The increased incidence of miscarriages in older women is likely due to the increased levels of MI nondisjunction (MI-ND; NAGAOKA *et al.* 2012).

Budding yeast has long served as an excellent model organism for studying the chromosome events of MI prophase. Forward genetic screens have identified dozens of conserved genes involved in key events such as homolog pairing, crossing over by homologous recombination, rapid chromosome motion and the formation of the monopolar attachment of sister chromatid pairs that insures proper disjunction between homologs (ZICKLER AND KLECKNER 2015). The ability to characterize these events in mutant strains has provided great insights into the functions of these individual proteins and their partners.

A defining phenotype of these meiotic mutants is the formation of inviable spores products. Spore inviability due to random spore death (RSD) can arise by the misappropriation of cellular components during spore-wall formation. Mitotic defects may also result in germination defects. Alternatively, defects in double strand break repair during meiosis may result in lethal lesions independent of segregation defects. More likely in the meiotic mutants described above, spore inviability is due to one or more spores of a tetrad failing to inherit one or more chromosomes (ROCKMILL *et al.* 2006). While every yeast chromosome is essential to support cell division, the presence extra chromosomes (n + 1) are generally viable (CAMPBELL AND DOOLITTLE 1987; TORRES *et al.* 2007; ST CHARLES *et al.* 2010).

The laboratory yeast strain, SK1, sporulates at high efficiency and with high levels of spore viability (∼98%). This feature makes it an excellent strain background for analyzing mutants defective for meiotic chromosome segregation and consequently spore death (MARTINI *et al.* 2006; WANAT *et al.* 2008). The proportion of live:dead spores in a tetrad gives some insight into the nature of the lethality. That is, a 3:1 live:dead tetrad would result from precocious sister chromatid separation (PSCS) at the meiosis I (MI) division or MII nondisjunction, while a 2:2 live:dead tetrad would result from MI-ND (Fig. 1; ROCKMILL *et al.* 2006). Events involving multiple chromosomes can give 1:3 or 0:4 live:dead spores. Thus, a good approximation of increased MI-ND can be inferred from a high incidence of 2:2 and 0:4 tetrads. Inviability is generally correlated with the severity of the mutant defect. Mutations that abolish homolog engagement may exhibit less than 1% spore viability while a mutation that disrupts chromosome movement during prophase may give only a modest reduction in spore viability (MARTINI *et al.* 2006; WANAT *et al.* 2008).

**Figure 1.**
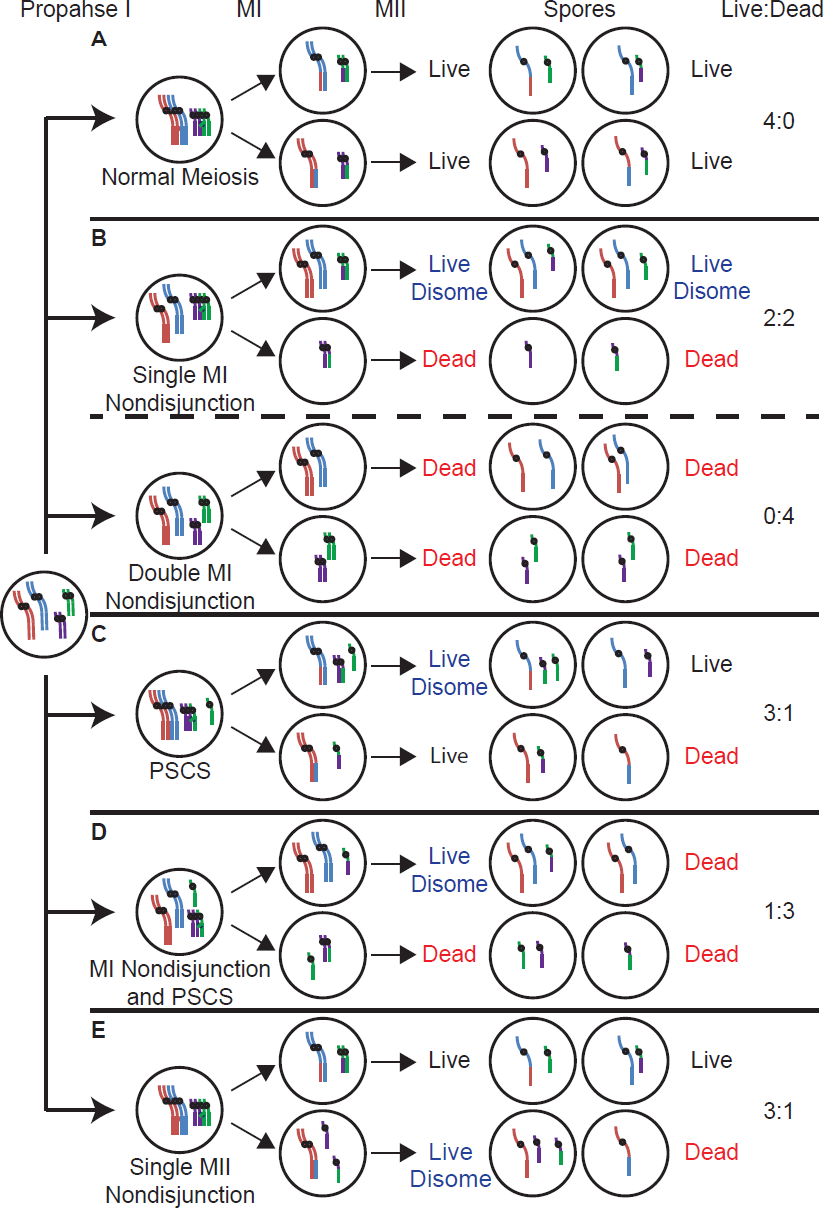
Schematic of different forms of nondisjunction. Newly replicated chromosomes (right) proceed to two sequential rounds of chromosome separation. The cell has two pairs of homologs, a long pair (red and blue) and a short pair (green and purple). Every chromosome is essential and the absence of either a long or short chromosome results in a dead spore. A spore with a single long and short chromosome is considered live and normal. A spore with either two long chromosomes and one short chromosome or one long chromosome and two short chromosomes is considered a live disomes. The live disomes have a high probability of living but may die at a low frequency due to being disomic. A) (Normal meiosis) The long and short homologous chromosomes each undergo crossing over to ensure their proper segregation at MI. Following MII, each spore receives a long and a short chromosome resulting in four live spores. B) (top; MI single-nondisjunction) The long pair of chromosomes fails to form a crossover resulting in an MI nondisjunction event where both chromosomes segregate to the same pole. Following MII, two spores are disomic for the long chromosome, and in most cases produce live spores. The two spores missing a copy of the long chromosome die. (Bottom; MI double-nondisjunction) Both the long and short chromosomes fail to form a crossover leading to a double nondisjunction where each pair of homologs separate to opposite poles. The result is four dead spores with two spores missing a long chromosome and two spores missing a short chromosome. C) (Precocious sister chromatid separation) A loss of sister chromid cohesion on a pair of short sister chromatids causes a sister chromatid to fail to properly disjoin during MI segregation. The sister chromatid segregates randomly during MI segregation resulting in two live spores, one live but disomic spore with an extra short chromosome, and one dead spore missing a short chromosome. D) (MI nondisjunction plus precocious sister chromatid separation). The long pair of chromosomes fail to form a crossover and there is a loss of sister chromatid cohesion between a pair of short sister chromatids. Only one spore is live but disomes and the rest of the spores are dead because they are either missing a long or short chromosome. E) (Single MII nondisjunction) Meiotic prophase and MI segregation are normal. Before MII segregation there is loss of sister chromatid cohesion on a pair of short sister chromatids causing them to segregate to the same pole. There are two live spores, one live but disomic spore with an extra short chromosome, and one dead spore missing a short chromosome.

Quantitative estimates of the frequency of MI-ND are typically carried out on a chromosome-by-chromosome basis and can require dissecting up to thousands of tetrads to generate accurate MI-ND frequencies. Such assays typically involve the detection of segregating heterozygous co-dominant genetic markers among the viable spore clones. Examples of this approach include co-segregation of *MATa* / *MATa* to form non-mating spore clones for chromosome III and the use of CEN-linked genetic markers (e.g. *URA3* / *TRP1)* for other chromosomes (WANAT *et al.* 2008). Alternatively, fluorescent methods can be used to detect nondisjunction in intact tetrads. One method is to tag specific chromosomes with large arrays of *tetO* or *lacO* DNA sequence (MARSTON *et al.* 2004). The chromosome can then be tracked by expressing either TetR or LacI fused to a fluorophore e.g. GFP, which will bind to the arrays allowing the presence of the tagged chromosomes to be visualized. Alternatively, a homolog pair can be engineered to express two different fluorophores from each homolog (THACKER *et al.* 2011). In the case of MI-ND, some spores will harbor multiple fluorophores while others will have none. Both of these cases have the advantage that many tetrads can be assayed without dissection, however, substantial strain construction carrying all relevant markers is first required.

Here we offer a computational method to quantify the average frequency of MI-ND per chromosome pair by determining the coefficient of nondisjunction that allows a best-fit distribution of 4:0, 3:1, 2:2, 1:3 and 0:4 live:dead spores to approximate the observed distribution. This method offers the advantage of scoring a MI-ND event involving of any of the 16 pairs of homologs requiring the dissection of fewer tetrads to assay MI-ND. We have compared our calculated rates to those determined empirically based on the mating type and marker segregation assays described above. Our results suggest that MI-ND is one of the main contributors to spore inviability in wild-type (WT) cells. These computational tools can be applied to any set of tetrad data to determine a calculated average nondisjunction frequency for any given strain and relative contribution of spore invability due to MI-ND.

## Methods

### Calculating the distribution of tetrad types based on random spore death

R-Script1 simulates the expected distribution of tetrads giving 4, 3, 2, 1, or 0 viable spores due to random spore death (RSD; File S1). Here we define RSD as the chance a spore will randomly die and that whether a spore lives or dies will not affect the behavior of other spores. In brief, the R-Script1 generates a matrix with the dimensions of 4 X number of tetrads, which is then filled with random numbers between 0 and 1. The random numbers are then converted to 1 (live) if the number is less than the spore viability set or 0 (dead) if the number is greater than the spore viability set. Each set of four spores generated are then converted to a hypothetical tetrad giving 4, 3, 2, 1, or 0 viable spores and the frequency of each tetrad class is calculated and recorded. This process is then repeated for the specified number of simulations and the distributions can be plotted using the Beeswarm package of R. For this paper 50,000 simulations were recorded every time the R-Script1 was used.

### Modeling the effects of RSD and MI nondisjunction on live:dead tetrad distributions

Using the assumptions of the effects of MI nondisjunction (MI-ND; discussed below), we created R-Script2 (File S2) which generates a matrix of live:dead tetrad distributions based on RSD and MI-ND values. Distributions of RSD and MI-ND set by the user are generated and then all pair wise combinations of RSD and MI-ND are added to the matrix. For each RSD and MI-ND pair, R-Script2 calculates the live:dead tetrad frequencies using the following equations:

First for each number of live spores possible (L, 0-4), R-Script2 calculates a binomial expansion to determine the live:dead tetrad frequencies (*f*L) based on the given RSD using the equation:

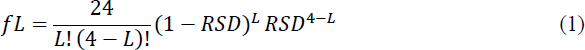

For each number of nondisjunction chromosomes (C, 0-16), the script next calculates the frequency of C chromosome nondisjunction events per meiosis (*f*C) using the MI nondisjunction rate (N) through a binomial expansion similar to the first equation:

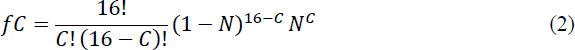

When C is greater than one, N is multiplied by 10 (the nondisjunction multiplier) to account for the nonrandom distribution of nondisjunction events. The nondisjunction multiplier was empirically determined as the value giving a best-fit live:dead tetrad distributions for the combined Shonn, Martini, and Wanat data sets as a test case. The aneuploidy induced death multiplier was similarly determined.

We then calculated the effects of aneuploidy-induced death for each C (ANID_C_) frequency by multiplying ANID and C. If ANID_C_ is greater than one then it is converted to one. The frequency of aneuploidy-induced death of 0, 1, and 2 spores per meiosis (*f*ANID0_C_, *f*ANID1_C_, and *f*ANID2_C_, respectively) for each C were calculated though a binomial expansion. To calculate the frequency of 2, 3, or 4 spores dead (*f*2SD_C_, *f*3SD_C_, and *f*4SD_C_, respectively) due to nondisjunction from C nondisjunction chromosomes and aneuploidy induced death (ANID) we used the following equations:

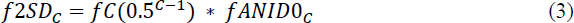

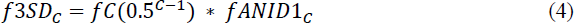

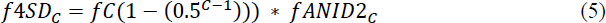

The *f*2SD_C_, *f*3SD_C_, and *f*4SD_C_ are then each summed for each C to generate the total frequencies of 2, 3, or 4 spores dead from all MI-ND events (*f2SD, f3SD,* and *f4SD*). The frequencies of spore death due to MI-ND are then applied to the previously calculated live:dead tetrad distributions calculated for RSD to generate the final live:dead tetrad distribution for a given MI-ND and RSD.

To calculate Avg-ND, R-Script2 uses the equation:

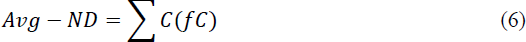

To calculate NDD, R-Script2 uses the equation:

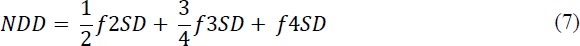

Although RSD and NDD can be directly compared to assess their relative contributions to spore inviability, they cannot be added together to determine spore viability because RSD and NDD will double count dead tetrads,

### Determining the best fit RSD and MI-ND

Once the all of the live:dead tetrad distributions have been calculated for each RSD and MI-ND, each generated live:dead tetrad distribution is compared to the observed distributions for each genotype using a chi-square test. The for each genotype, the live:dead tetrad distribution with the highest P value is recorded as the best fitting tetrad. P values were adjusted using the Holm method (HOLM 1979).

## Results

### RSD alone does not account for the observed distribution of live:dead tetrads from WT strains

WT strains of budding yeast typically give around 98% spore viability among tetrads (Fig. 2). We wondered whether or not the 2% spore inviability in WT cells could be attributed solely to RSD. To test this we created R-Script1 to model the expected distribution of live:dead tetrads due to RSD based on empirically measured spore viability frequencies. These calculated distributions were then compared to a set of six empirically measured data from WT strains as reported in six previously published papers (MASISON AND BAKER 1992; SHONN *et al.* 2000; JESSOP *et al.* 2006; MARTINI *et al.* 2006; WANAT *et al.* 2008; KEELAGHER *et al.* 2011).

**Figure 2.**
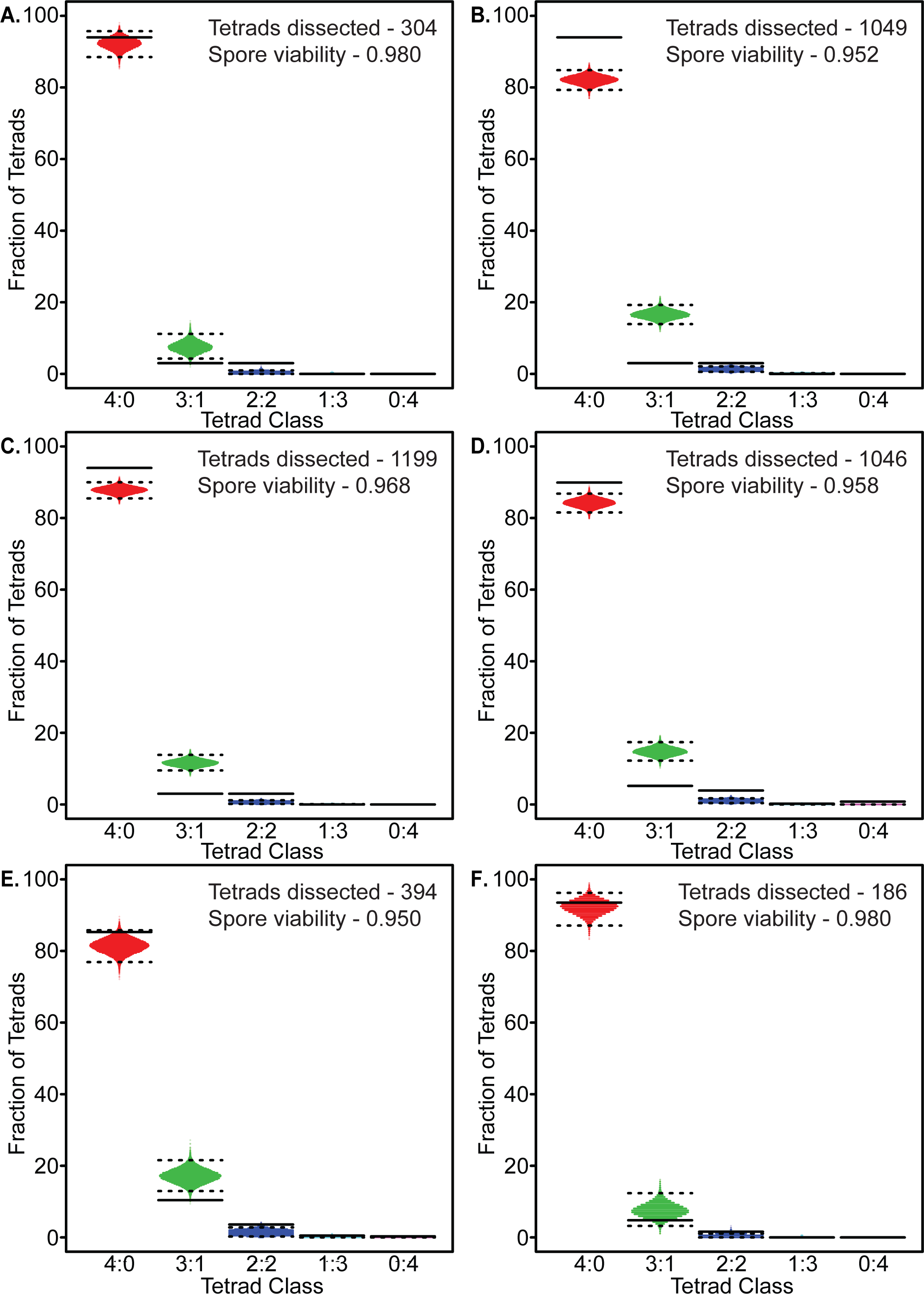
Simulation of random spore death. Representative simulation of tetrad distributions using R-Script1 based observed spore viability and number of tetrads dissected for the indicated data sets. A Poisson distribution of dissected tetrad simulations is shown with the upper and lower dashed lines represent 99% and 1% percentiles, respectively. The observed frequencies of live:dead tetrad distributions for individual strains are shown as solid lines. A-F Comparison of simulated tetrad distributions with WT data from previously published data sets. A) Shonn et al; B) Martini et al; C) Wanat et al;. D) Keelagher et al; E) Masison et al; F) Jessop et al.

If the spore inviability in the six WT strains was solely due to RSD, then the observed distribution of live:dead tetrads should resemble the output of R-Script1. We found, however, that the observed distributions reported for all WT strains fell outside of the 1 and 99 percentiles of simulated data (Fig. 2). Specifically, the published data showed an overrepresentation of 2:2 live:dead tetrads and underrepresentation of 3:1 live:dead tetrad distributions compared to the simulation (Fig. 2). The observed pattern could not be modeled by simply increasing or decreasing the simulated spore viability. That is, increasing the simulated spore viability would consequently decrease the frequency of 2:2 live:dead tetrads. Conversely, decreasing the simulated spore viability increased the frequency of 3:1 live:dead tetrads. These results suggest that spore inviability in WT strains is not solely due to RSD. Instead, the increased incidence of 2:2 live:dead tetrads observed for WT compared to the simulation suggests that spore death may be due to an intrinsic level of MI-ND.

### Modeling MI-ND

We next tested whether or not the observed increased levels of 2:2 live:dead tetrads for both WT and mutant strains reported in the published data sets could be attributed to MI-ND or a combination of MIND and RSD events. We created R-Script2 to model the relative contributions of RSD and MI-ND events that give the best fit to the observed data. Script2 also generates the average MI nondisjunction (Avg-ND) frequency, which is the frequency of any given chromosome undergoing a MI-ND event per meiosis. The Avg-ND frequency should be comparable to previously published MI-ND frequencies for individually-tested chromosomes (Table 1).

**Table 1.**
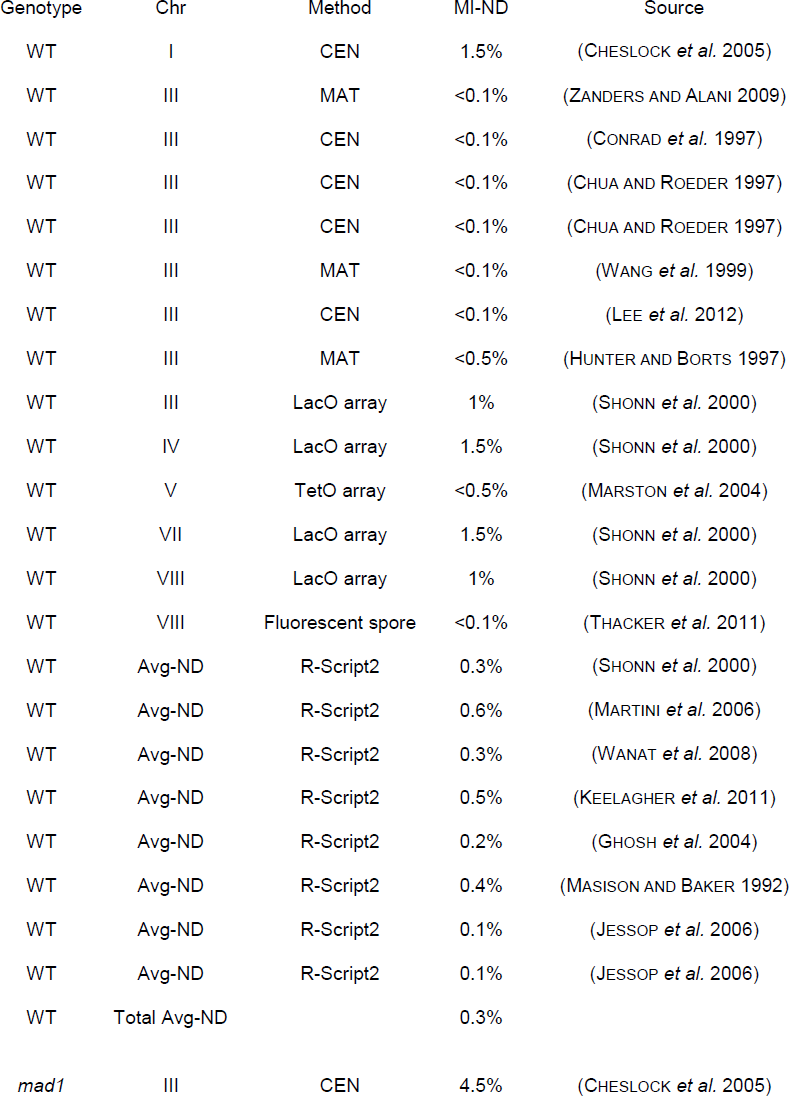
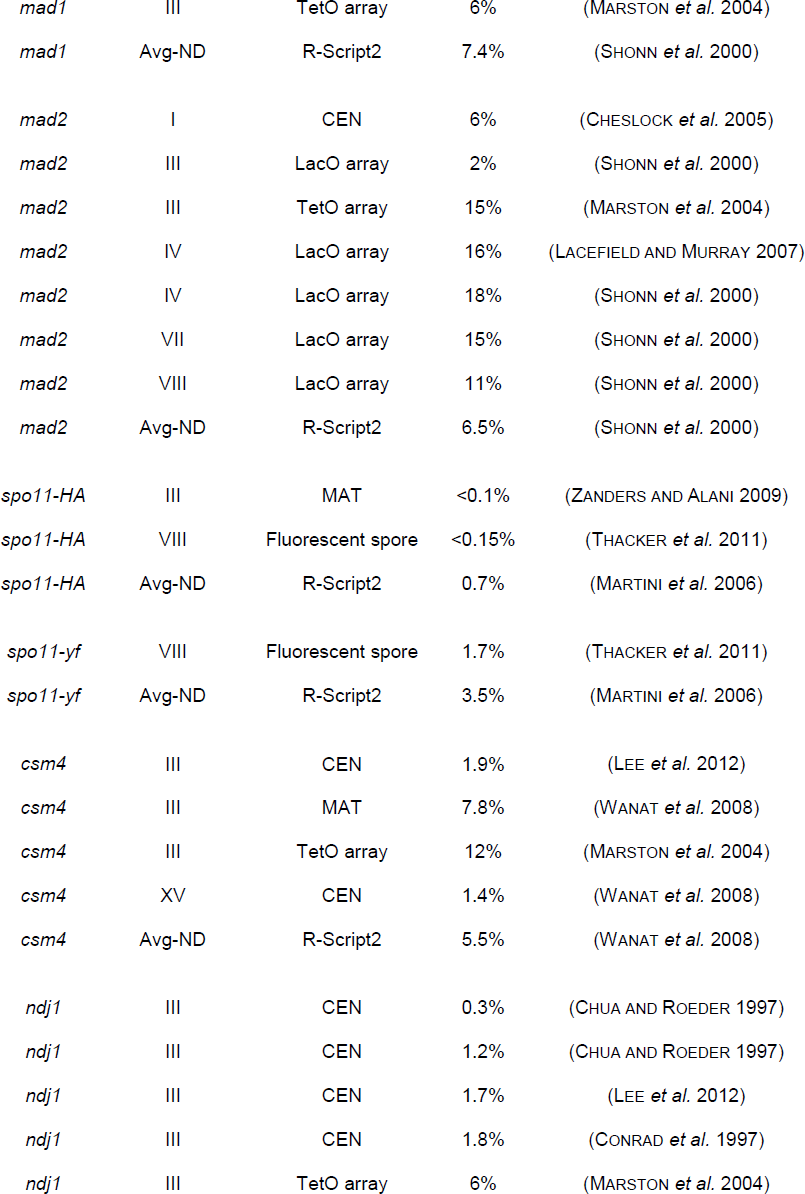
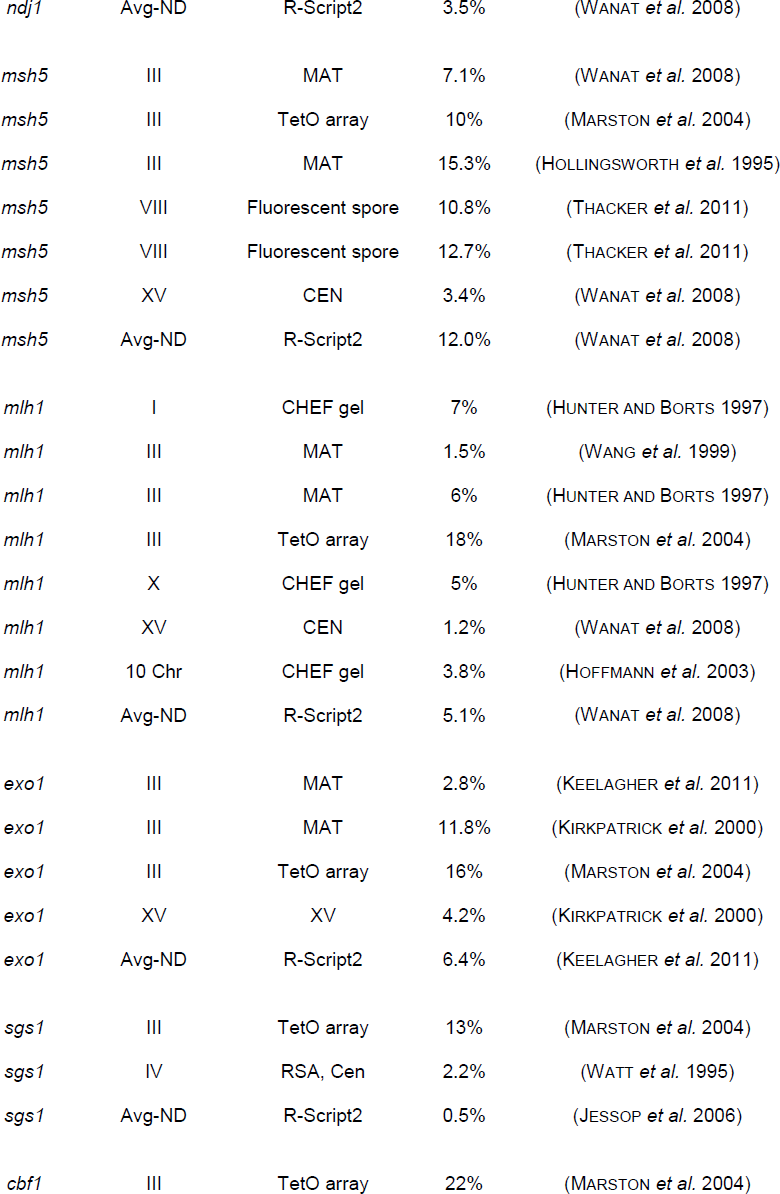
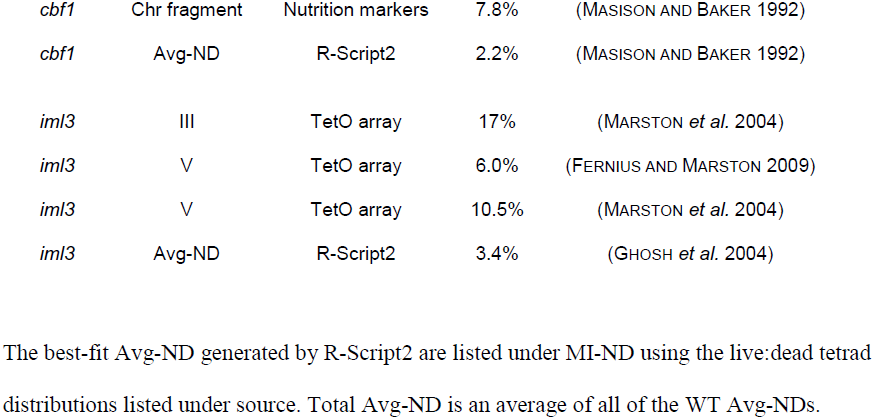
**Empirically derived MI-ND frequencies and computationally generated Avg-ND frequencies.**

R-Script2 takes into account several features of chromosome aneuploidy. First, MI-ND is defined in the classical sense in which two homologs fail to properly disjoin and segregate to the same pole (Fig. 1B). Here we assumed that every chromosome is essential and that the MI-ND frequency for each chromosome is equal. As most disomies are well tolerated in yeast, most tetrads which have undergone a MI-ND event will give 2 live and 2 dead spores (TORRES *et al.* 2007; ST CHARLES *et al.* 2010). In the case of multiple MI-ND events, we assumed that the homologs would segregate randomly to each pole. When both pairs of homologs segregate to opposite poles, all four spores would be inviable. Conversely, if both homolog pairs were to separate to the same pole, then two of the spores would live but doubly disomic and the other two spores would be dead.

Second, the incidence of multiple MI-ND events does not follow a Poisson distribution (SHONN *et al.* 2000); i.e., there are more multi-chromosome MI-ND events than expected from a random distribution. To account for this increase, we used an empirically determined multiplier of 10 across all data sets in the instances of multiple MI-ND events based on the live:dead tetrad distributions reported by Shonn et al. Martini et al., and Wanat et al.

Third, the effects of multiple disomies appear to be relatively low since spore viability rates from triploid meiosis, where nearly every spore would contain multiple disomic chromosomes, can be as high as 50-83% (CAMPBELL AND DOOLITTLE 1987; ST CHARLES *et al.* 2010). Disomies of specific chromosomes, however, have been shown to reduce cell fitness and germination rates (CAMPBELL AND DOOLITTLE 1987; TORRES *et al.* 2007; ST CHARLES *et al.* 2010). To account for negative effect of multiple disomies on spore germination, we included an empirically-determined frequency of aneuploidy-induced death (3.5% multiplied by the number of disomic chromosomes) based on the three published data sets.

### Proof of concept

We selected a battery of mutants known to increase either MI-ND or PSCS/MII nondisjunction. Since PSCS and MII-nondisjunction events would appear as RSD, we reasoned that mutants that give high incidence of PCSC and MII-ND would be expected to give relatively high levels of RSD. We then applied R-Script2 to generate distributions of live:dead tetrads to find the best-fit distribution to each observed data set. The mutants with known defects in MI-ND included *mad1Δ* and *mad2Δ* mutants that abrogate the spindle assembly checkpoint (SHONN *et al.* 2000), the hypomorphic alleles of *SPO11* that reduce the levels of DSBs (*spo11-HA*, *spo11-yf,* and *spo11-df* (MARTINI *et al.* 2006), *ndj1Δ* and *csm4Δ* mutants defective for telomere-led motion, and *msh5Δ*, *mlh1Δ*, and *exo1Δ* mutants that affect homologous recombination (WANAT *et al.* 2008; KEELAGHER *et al.* 2011). We also included in our analysis *cbf1Δ* and *iml3Δ* mutants with defective centromeres (MASISON AND BAKER 1992; GHOSH *et al.* 2004), and *sgs1Δ* and *sgsΔ795* mutants that display increased PSCS due to increased crossover formation near centromeres (JESSOP *et al.* 2006).

### Modeled tetrad distributions closely match observed tetrad distributions in WT and mutant strains

A number of mutants have been shown to give relatively increased levels of 2:2 and 0:4 live:dead tetrads. We focused on data sets from studies where the distribution of live:dead tetrads and the nondisjunction frequency of one or more test chromosomes were analyzed (MASISON AND BAKER 1992; SHONN *et al.* 2000; GHOSH *et al.* 2004; JESSOP *et al.* 2006; MARTINI *et al.* 2006; WANAT *et al.* 2008; KEELAGHER *et al.* 2011). We applied R-Script2A to find the expected (E) RSD and ND frequencies that give the best fit to the observed (O) tetrad distributions for each mutant (Fig. 3). We also applied R-Script2B to find the best fitting tetrad distribution and RSD when using each mutant’s respective WT (W) MI nondisjunction rate (Fig. 3).

**Figure 3.**
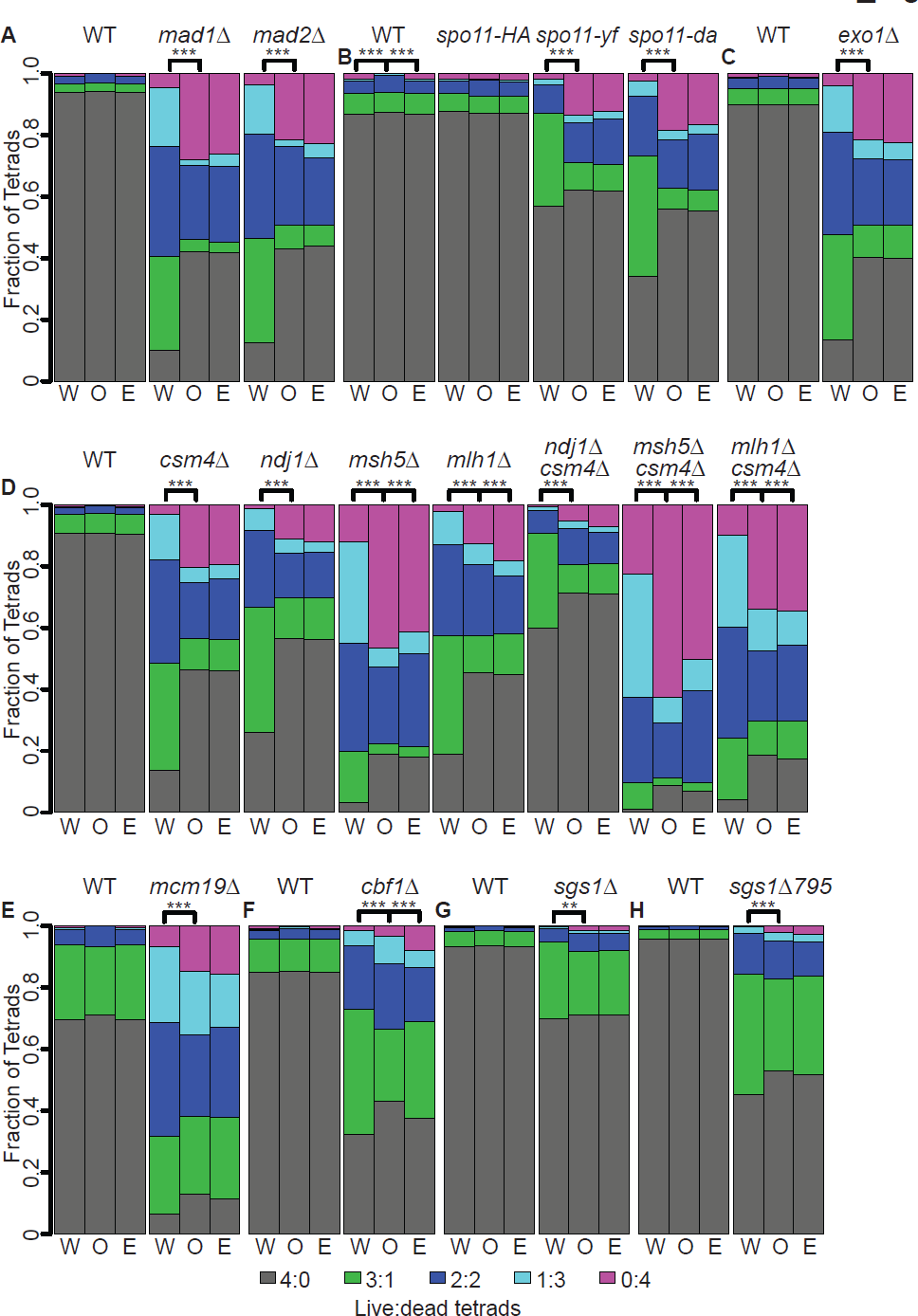
Comparison of observed and expected tetrad distributions. The observed tetrad distributions of genotype (O, middle bar), the expected tetrad distributions with the best fitting MI-ND and RSD (E, right bar), and the expected tetrad distributions using WT MI-ND and the best fitting RSD (W, left bar). Expected live:dead tetrad frequencies were generated for all genotypes using R-Script2 with the following conditions: the number of nondisjunction intervals (ndint) was set to 3000, the number of random spore death intervals (rsdint) was set to 3000, the number of chromosomes (chr) was set to 16, the aneuploidy induced death frequency (anid) was set to 0.035, and the nondisjunction multiplier (ndm) was set to 10. (A-G) Comparison of simulated tetrad distributions with WT and mutant data from previously published data sets: A) Shonn data set; B) Martini data set; C) Keelagher data set; D) Wanat data set; E) Ghosh data set; F) Masison data set; G) Jessop data set; H) Jessop data set. In these strains the endogenous SGS1 has been deleted but *SGS1* or *sgs1Δ795* have been inserted at TRP1 in the WT and *sgs1Δ795* strains, respectively. Significance is noted as **P < 0.005; ***P <0.0001.

For the MI-ND mutants we found that R-Script2 generated live:dead distributions that were statistically indistinguishable from all of the mutant distributions with the exceptions *msh5Δ* and *mlh1Δ* (chi-square test, Holm correction; Fig. 3 D). Msh5 and Mlh1 play the most direct role in the mechanism of DSB repair among the MI-ND set mutants. One possible explanation for the poor fit is that the low sporulation efficiency in the mutants may lead to biases in the distributions (BORNER *et al.* 2004). Alternatively, segregation of unrepaired recombination intermediates in these strains could introduce a variable that is not accounted for by R-Script2.

The observed and expected distributions for the *cbf1*Δ strain also failed to show a significant difference from the distribution using WT nondisjunction levels. Loss of Cbf1 has be shown previously to increases PSCS frequency (Fig. 3F; MASISON AND BAKER 1992). Since Cbf1is a transcription factor, its absence may cause multiple defects in addition to its role in preventing PSCS (BAKER AND MASISON 1990; MASISON AND BAKER 1992). Moreover, analysis of *cbf1*Δ was carried out in a 381G background strain, which was different than the other strains analyzed (MASISON AND BAKER 1992). Despite these differences, the overall expected and observed distributions are qualitatively similar.

All the mutants we looked at gave distributions that differed significantly from a model based soley on RSD and using each mutants respective “WT” MI-ND frequency (W; Fig. 3). The one exception was *spo11-HA,* which was not surprising since it gives WT levels of spore viability (MARTINI *et al.* 2006). Together these results show that R-Script2 gives an accurate estimate of MI-ND frequencies in WT and mutant strains.

### Comparison of Avg-ND to observed ND for individual chromosomes

Estimates of MI-ND frequencies can vary widely depending on the assay and chromosome analyzed (Table 1). For example, for the *mad2Δ* mutant measurements using the same assay and the same strain background gave MI-ND frequencies ranging from 2%-18% for different chromosomes (Table 1; SHONN *et al.* 2000). R-Script2 calculates average nondisjunction rates for all chromosomes, thus eliminating the risk of picking a chromosome with much higher or lower level of MI-ND compared to other chromosomes. Conversely, it is important to consider that the Avg-ND frequencies generated by R-Script2 represent an average for all of the chromosomes, and does not provide information on an individual chromosome.

Thus we expected that the Avg-ND frequency for WT and the mutants affecting MI nondisjunction would fall within the range of observed frequencies for individual chromosomes. For the mutants described above that were shown previously to increase MI chromosome nondisjunction, the Avg-ND was within the range of measured values, if not slightly higher (Table 1). By contrast, for the mutants shown previously to increase PSCS, the calculated Avg-ND tended to be slightly lower (Table 1). This could be due to the fact that a subset of PSCS events will be counted as a nondisjunction event in the given assays, while the calculated Avg-ND value does not include PSCS events. These results indicate that Avg-ND gives an approximate value of nondisjunction thus allowing direct comparisons between mutant strains.

### The predictive power of a computational approach to measuring nondisjunction frequency

MI-ND and RSD (including PSCS and MII-ND events) contribute independently to spore inviability (above). To directly compare the contributions of death from MI nondisjunction death and RSD, we computationally converted the expression of MI-ND frequency to nondisjunction death (NDD). For WT strains the contribution of NDD and RSD were roughly equal (Fig. 4). Mutant strains appeared to fall into two classes: The first class (Class I) gave NDD/RSD > 3.3 while the second class (Class II) gave NDD/RSD < 0.8 (Fig. 4). Not suprisingly, Class I genes include those with roles in homolog engagement and separation while Class II genes include roles in sister chromatid cohesion and centromere function.

**Figure 4.**
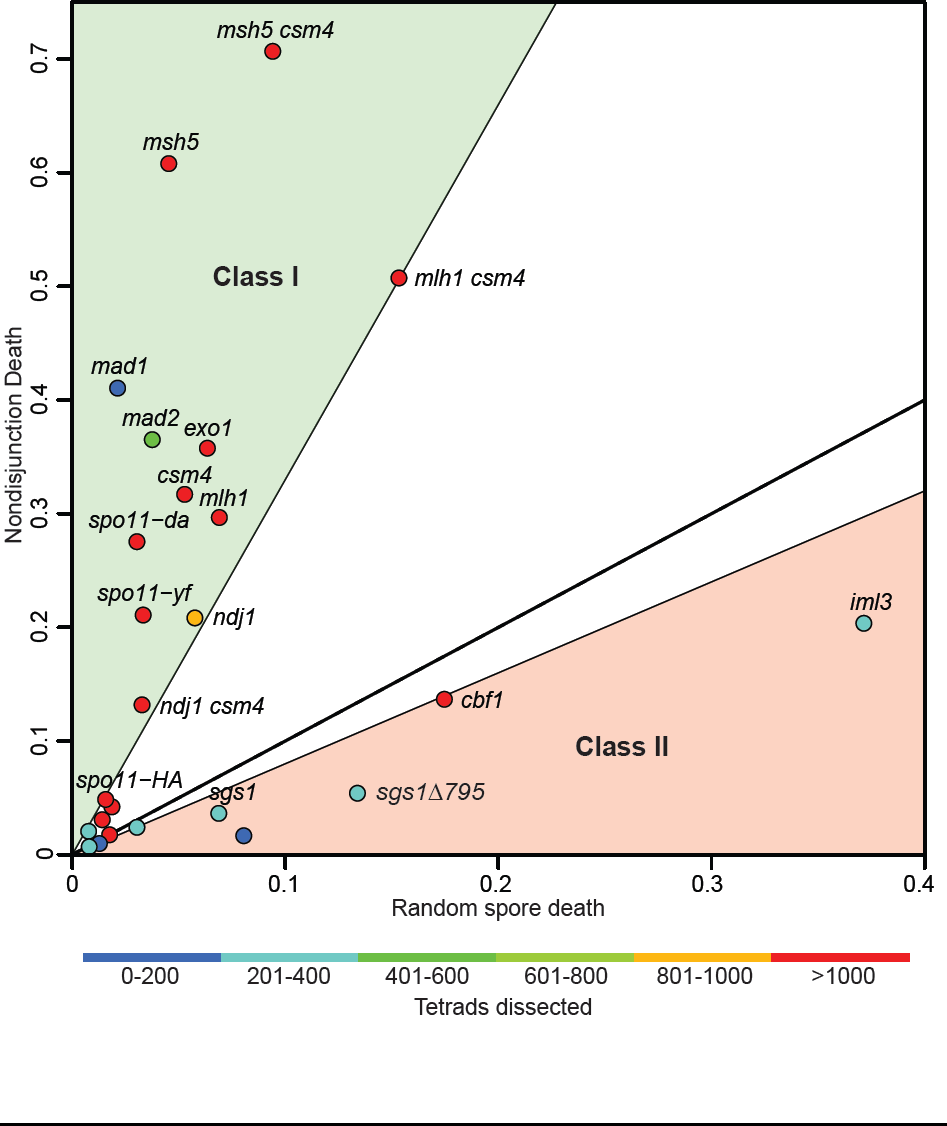
Comparison of random spore death and nondisjunction death. A scatter plot of the calculated best fits of RSD and NDD for all of the data sets. All WT strains are shown as unlabeled spots on the plot. Data points are colored based on the reported number of tetrads dissected. The solid line represents a NDD/RSD slope of one which indicates an equal contribution of NDD and RSD on spore inviability. The green shaded area covers the Class I mutants which have an NDD/RSD ratio above 3.3 and the red shaded area covers Class II mutants which have an NDD/RSD ratio below 0.8.

The ability to separate the relative contributions of RSD and NDD is also a useful tool for genetic epistasis analysis. Both the *csm4Δ msh5Δ* and *csm4Δ mlh1Δ* double mutants give increased levels of RSD and NDD compared to their respective single mutants (Fig. 5A). The NDD/RDS ratio for *csm4Δ msh5Δ* is intermediate to those for each single mutant, suggesting that *csm4Δ* and *msh5Δ* both increase NDD and RSD through independent pathways.

**Figure 5.**
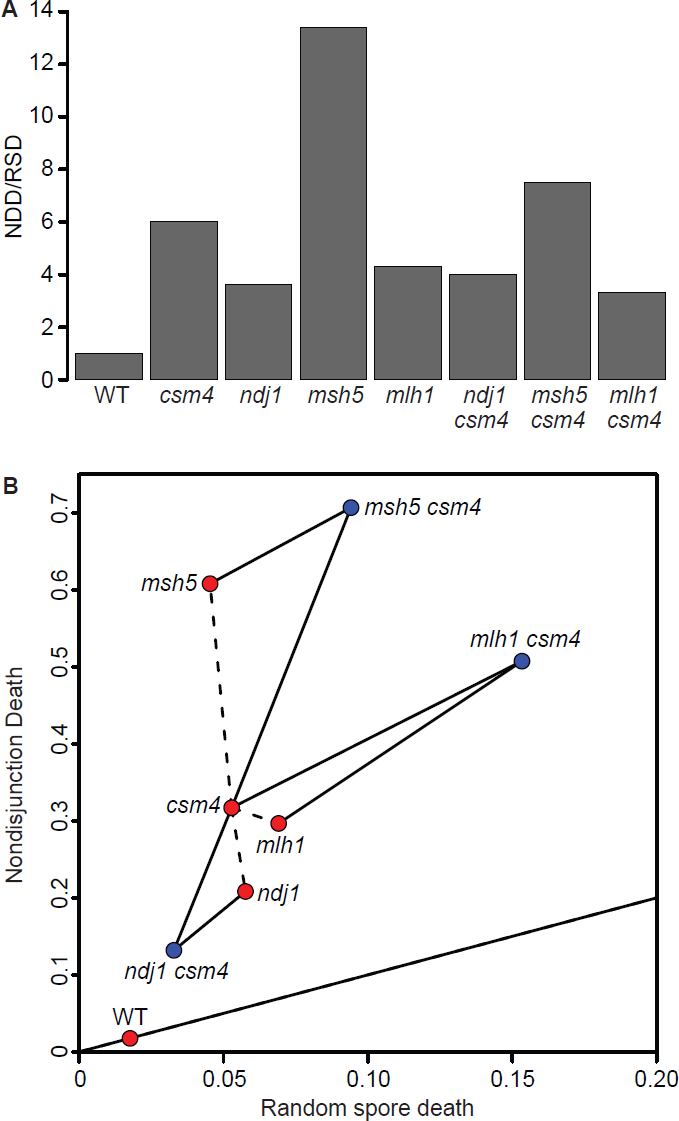
Genetic analysis of double mutants. A) Bar plot of the NDD/RSD ratios from the Wanat data set. B) Scatter plot of RSD and NDD using the Wanat data set. Single mutants and WT are shown in red and double mutants are shown in blue. Solid lines connect the double mutants to their respective single mutants and dashed lines connect the single mutants together. A solid line with a NDD/RSD ratio of one is shown.

By contrast, the NDD/RSD ratio for *csm4Δ mlh1Δ* is lower than for each single mutant, which could indicate less NDD and/or more RSD than expected for an additive interaction. By comparing the absolute NDD and RSD values it appears that *csm4Δ mlh1Δ* exhibits less NDD and more RSD than expected, suggesting a more complex relationship between the two mutants than previously observed (WANAT *et al.* 2008). That is, the two mutations act together to reduce NDD, perhaps at the expense of increased RSD. Both RSD and NDD are reduced in the *ndj1Δ csm4Δ* double mutant both compared to either single mutant. The *ndj1Δ csm4Δ* double mutant has been shown previously to interact genetically in a partial-reciprocal epistastic manner in which the phenotype of the double mutant is weaker than either of the single mutants (WANAT *et al.* 2008). Our results are consistent with such a relationship since it appears that is that *csm4Δ* suppresses the increased PSCS phenotype of the *ndj1Δ* mutant and that *ndj1Δ* suppresses the NDD phenotype of *csm4Δ*.

## Discussion

In this study, we created R-Script1 to test if an observed live:dead tetrad distribution fits an expected distribution based on RSD. We found that spore death in WT strains could not be accounted for by RSD alone. We next created R-Script2 to estimate the relative contributions of NDD and RSD on final spore viability and live:dead tetrad distributions. The script has advantages over other methods of nondisjunction estimation by using the information from all dissected tetrads including 0:4 live:dead tetrads. Additionally R-Script-2 analyzes the nondisjunction of all chromosomes holistically instead of analyzing single chromosomes.

Considering all potential chromosome nondisjunction events has the advantage over a single chromosome approach since estimated nondisjunction rates can be determined from a much smaller sample size. For example, for the *csm4Δ* mutant, only 3 tetrads in 1164 total dissected had a chromosome XV nondisjunction event detectable by CEN markers (WANAT *et al.* 2008). To generate more accurate estimations of nondisjunction would require many more labor-intensive tetrad dissections. If all chromosomes were to have an equal chance of nondisjunction and only 1 out of 16 chromosomes is analyzed, only ∼6% of an already rare event can be detected. In the case of multiple nondisjunction events the resulting 0:4 live:dead tetrads (Fig. 1B) could not be analyzed. This would also lead to biased sampling since only a fraction of nondisjunction events involving the test chromosome would be detected. Because of these reasons several thousands of tetrads would need to be dissected to detect a few rare nondisjunction events for only a subset of chromosomes.

Using R-Script2 described here requires the analysis of only 100 -200 dissected tetrads to generate RSD and MI-ND data for analysis. The low numbers of tetrads required for each strain also facilitates epistasis analysis. Here we show as proof-of-principle that the analysis of single and double mutant strains can be performed with few tetrads. Our analysis also provides additional insight into the nature of spore death that occurs in the double mutants versus single mutants. Importantly, the application of the R-Scripts does not require any special strain construction and can be applied to previously observed tetrad distributions.

